# Event-Related Desynchronization induced by Tactile Imagery: an EEG Study

**DOI:** 10.1101/2022.11.09.514848

**Authors:** Lev Yakovlev, Nikolay Syrov, Andrei Miroshnikov, Mikhail Lebedev, Alexander Kaplan

## Abstract

It is well known that both the movement of the hand itself and the mental representation of it lead to event-related desynchronization (ERD) of EEG recorded over the corresponding motor areas of the cerebral cortex. Similarly, in somatosensory cortical areas, ERD occurs upon tactile stimulation of the hand, but whether this effect is caused by mental representation of sensations from tactile stimulation remains poorly understood. In the present study, the effects on the EEG of imaginary vibrotactile sensations on the right hand were compared with the effects of real vibrotactile stimulation. Both actual vibrotactile stimulation and mental representation of it have been found to elicit contralateral ERD patterns, particularly prominent in the *μ*-band and most pronounced in the C3 region. The paper discusses tactile imagery as a part of the complex sensorimotor mental image and its prospects for using EEG patterns of imagery-induced tactile sensations as control signals in BCI circuits independently and when combined with ERD based on movement imagination to improve the efficiency of neurointerface technologies in rehabilitation medicine, in particular, to restore movements after a stroke and neurotrauma.

## INTRODUCTION

The ability to construct inner mental representations of real external activities (mental imagery) is a fundamental aspect of human cognition. Mental imagery is associated with such comprehension-requiring functions as memory, creativity, motor control, navigation, arithmetic, moral decisions and mind wandering [1].

One of the most studied mental imagery processes is related to movements. It is known as motor imagery (MI). It has been shown that motor imagery induces neuroplasticity changes at the cortical level and facilitates motor learning [2, 3]. These effects allow use of MI practice as a supplementary physical training in sports [4], as well as a form of neurorehabilitation for functional recovery after strokes [5]. Depending on attentional focus associated with the imagined movement, kinesthetic and visual types of motor imagery are distinguished [6]. These two distinct strategies have specific effects on corticospinal excitability [7], EEG activation patterns [8] and metabolic brain activation [9]. These studies suggest that kinesthetic motor imagery is processed by the motor circuits that overlap with the circuits activated during the preparation and execution of voluntary movements. Additionally, since imagining sensations arising from muscle contraction (muscle and skin tension, changes in joint angle) is typically a part of kinesthetic cognitive imagery strategy, somatosensory processing is involved, as well. The somatosensory component of motor imagery was studied experimentally [10] and analyzed theoretically [11]. Thus, Natio et al. demonstrated that imagined kinesthetic sensations are associated with activity in multiple sensorimotor areas. Forukas, Ionta & Aglioti showed that imagined posture impacts cortical excitability [12]. Mizuguchi and his colleagues reported that motor imagery effects on corticospinal excitability were facilitated by somatosensory inputs generated by touching an object [13-15]. Additional evidence of influence of somatosensory inputs on motor imagery comes from the studies of neuromuscular electrical stimulation. Kaneko et al. reported that electrical stimulation of the index finger increases cortical excitability measured with single-pulse TMS [16]. We previously obtained similar results using functional electrical stimulation applied to the right hand while subjects imagined the hand moving [17].

Event related desynchronization (ERD), which is a transient sustained decrease in EEG/MEG power in a specific frequency band [18, 19], is a conventional marker of the presence of sensorimotor processing. ERD and the opposite process called event related synchronization (ERS) represent changes in correlated activity of the underlying neuronal populations, and these changes correspond to increased and decreased cortical processing, respectively [18]. As such, ERD and ERS can be used for functional cortical mapping [20, 21] or as a classification feature for brain computer interfaces (BCIs) [22-24]. In motor tasks, sensorimotor ERD occurs during voluntary movements, movements observation and motor imagery and in association with somatosensory inputs like tactile stimulation. Oscillatory activity over cortical sensorimotor regions is called somatosensory (or somatomotor) activity. This oscillatory activity comprises oscillations in the classic α (8-12Hz) and *β* (13-30) ranges [25]. These components have different sources. The sensorimotor α is generated in the primary somatosensory cortex, whereas sensorimotor *β* is generated in the primary motor cortex [25]. The involvement of primary somatosensory in the generation of *μ*-rhythm was confirmed by Frolov and his colleagues. They observed that the sources of motor imagery-associated EEG activity were localized in the hand representation of the sensorimotor cortex [26], and the BOLD signal had the same source [27].

While studies on motor imagery are plentiful, less work was conducted on tactile imagery. Neuroimaging studies of tactile imagery are limited to the fMRI method [28-31], and there is no systematic work with the EEG approach. Yoo et al. reported activation of the sensorimotor cortex caused by mental imagery of brushing stimuli [28]. Schmidt and his colleagues provided evidence of changes in coupling between cortical areas during tactile imagery [29, 31]. Furthermore, de Borst and de Gelder conducted an fMRI study of the activation patterns in visual, auditory, and somatosensory domains during imagery and actual perception [30]. Overall, these studies demonstrated similarity between neural patterns exhibited during real perceptions and tactile imagery tasks, which is consistent with the results of motor imagery studies showing that overlapping brain regions are engaged in actual movements and motor imagery. Additionally, in work [32] authors observed cortical activity in human subjects imagining the sensations evoked by intracortical microstimulation (ICMS) of their somatosensory cortex using an implanted microelectrode array. Thus, previous results on tactile imagery suggest that activation of somatosensory areas is a mechanism underlying tactile imagery.

Building on these previous results, here we sought for an EEG correlate of tactile imagery. By analogy with motor imagery, we hypothesized that tactile imagery would lead to modulations of the *μ*-rhythm, presumably in the contralateral hemisphere. We indeed discovered *μ*-rhythm ERD associated with both tactile imagery and real tactile stimuli – the findings that add to the literature on mental imagery and contribute to the development of new generations of BCI technologies.

## MATERIAL AND METHODS

### Participants

18 healthy right handed volunteers (9 females, age 22,5 ± 2,8) without a history of neurological disorders took part in this study. All of them had no prior experience in mental imagery of the motor and tactile types. The study protocol was approved by the Lomonosov Moscow State University ethical committee and followed the Declaration of Helsinki. All participants were informed about the experimental procedures and signed an informed consent.

### Experimental Procedure

The experiment duration did not exceed 90 minutes. During an experimental session, subjects sat in a comfortable armchair in the room with uniform lighting. The visual cues corresponding to experimental conditions were presented on a 24-inch LCD monitor positioned in front of the subject ∼1 meter away from their eyes.

The experiment included four different conditions: tactile stimulation (TS), learning of tactile imagery, tactile imagery (TI) and control condition. The experimental sequence is presented in Fig.1. Each experimental condition was organized as a sequence of randomly mixed trials with the duration of 6s. The trials represented two types of somatosensory tasks (TS or TI) or the visual attention task, which was used as a reference state to the somatosensory tasks. During the visual attention trials, participants observed an image with geometric elements: circles, dots, lines, and squares etc. (see Fig.1). Participants were instructed to keep their visual attention on the image and count all elements mentally avoiding explicit eye movements. Similar complex visual scenes were used as reference states in previous motor imagery studies [24, 33, 34]. The somatosensory trials were cued by a pictogram that instructed TS or TI. The gray screen was shown during the interval between trials, which lasted 300-400 ms.

**Fig 1.**
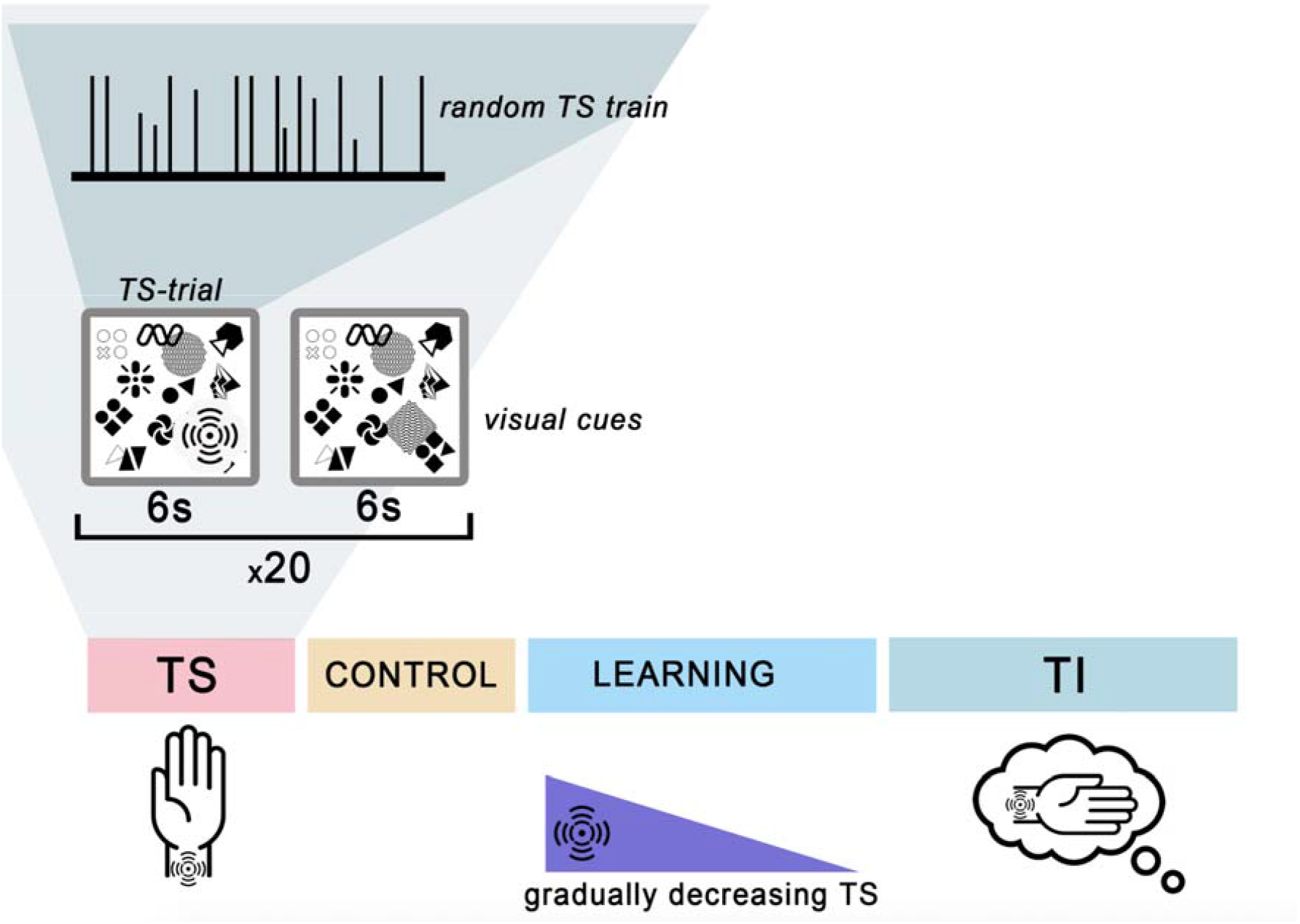
Block scheme of the experimental session. Each condition included 20 somatosensory trials and 20 visual-attention trials mixed randomly. In the TS trials the vibration intensity of the 6-s random stimulation train varied in range 3700-12000 rpm, and the time interval between the single vibrations varied in range 200-50 ms. Control trials were cued with the same visual stimulus but without any TS or TI performance. During TI trials no actual stimulation was performed.

**Table 1.** Detailed description of the experimental session.

| condition | runs | trials (in one run) | active trials, total |
| --- | --- | --- | --- |
| Tactile stimulation | 1 | 20 TS + 20 rst | 20 |
| Control | 1 | 20 ctrl + 20 rst | 20 |
| Learning | 2 | 20 TS/TI + 20 rst | 40 |
| Tactile imagery | 2 | 20 TI + 20 rst | 40 |

### Selection of the Site for Tactile Stimulation and Imagery

The skin area for the application of tactile stimulation was selected based on the tests in ten subjects, which preceded the main series of experiments. We sought for the place on the arm where tactile stimulation would be well perceived without any unpleasant sensations and could be reproduced in mental imagery. The subjects did not have any prior experience in imagery, and 4 of them participated later in the main experimental sessions. Vibrotactile stimuli were delivered with vibration motors. The motors were placed over several locations on the right arm, including index finger, thumb, palm and inner wrist surface. We used 20 trials per stimulation site, followed by the attempts to imagine the tactile sensation. The participants were then asked to describe their subjective experience and select the site where the stimulation-evoked sensation was the easiest to imagine. The subjects selected the most vivid and steady mental image.

Based on the results of these tests, the surface of the distal phalanx of the index finger and inner surface of the wrist were chosen as the most suitable places for TS and TI. We preferred this site and chose not to apply TS to the fingers for several reasons. First, finger stimulation led to long-term residual sensations that could continue into the other trials. Second, EEG modulations during finger stimulation are not hemispherically specific. According to [35, 36], mental imagery of finger movements, as well as tactile stimulation of the fingers, result in bilateral *μ*-ERD patterns whereas stimulation of more proximal parts of the arm cause predominantly contralateral ERD patterns [37, 38].

### Tactile Stimulation Technique

We used a custom designed and computer-controlled vibrotactile stimulator based on an Arduino UNO (Arduino, Spain). A flat vibration motor (6 mm in diameter, max speed of 12,000 rpm) was placed on the inner wrist surface of the right hand. The stimulation lasted throughout the 6-s trial. The stimulation pattern was chaotic: the motor speed varied in range 3700-12000 rpm, the one-stimulus duration was kept at 100 ms, and the time interval between the single vibrations varied in range 200-50 ms (Fig. 1). Such a random stimulation pattern reduces tactile habituation and helped to avoid residual tactile sensations. TS trials (N=20) were randomly intermixed with visual-attention trials (N=20).

### Transition to Tactile Imagery

Transition to TI was conducted gradually during the learning session that included 20 TS trials with decreasing stimulation amplitude and 20 visual-attention trials. The TS intensity decreased gradually from the parameters used in the TS session to zero. The subjects were instructed to focus attention on the skin sensations evoked by TS and to memorize them. As the TS intensity decreased, participants were asked to mentally compensate for the lack of stimulation by imagining the previously experienced sensation of vibration. A similar learning strategy was previously used in motor-imagery studies [24, 39-41].

Following the learning session, subjects provided a verbal report regarding task difficulty; they also described the imagery vividness. An objective measure of the imagery strength was given by the *μ*-ERD value and topographic distribution. The imagery was considered stable if a participant reported that he/she could mentally reproduce the tactile sensations at the same place where TS was applied, the imagined sensation was not associated with the limb movements and muscle contractions, and the imagery was maintained for 6s. The data from three participants were rejected from the analysis because one of them could not imagine tactile sensations and in the other two no ERD occurred during both TS and TI.

### Tactile Imagery

The TI trials began after the learning session. No tactile stimuli were applied, and participants were instructed to imagine vibrotactile stimulation being applied. These trials were triggered with the same cues as in the TS trials. The TI condition included two runs with a total number of TI trials of 40. TI trials were randomly intermixed with visual-attention trials. Participants were asked to maintain imagery of the randomly patterned vibrotactile stimulation for the entire 6s during which the cue was shown on the screen. We controlled for the absence of muscle activity using an online monitoring of the EMG signals from the right FDS muscle.

### Control Trials

Since the TS and TI trials were cued with the same visual stimulus (vibration pictogram), we had to control for the possible association of tactile sensations with this cue. Indeed, repeated presentation of such stimuli leads to the development of cross-modal association [42, 43]. As a control, we added the condition where participants observed the same visual cues as in TS and TI trials, but no stimulation or imagery happened. These trials were interspersed with the visual-attention trials. This control session includes 20 trials with the vibration-related cues and 20 visual-attention trials.

### EEG and EMG Recordings

We recorded 48-channel monopolar EEG data at 500 Hz sampling rate using an NVX-52 DC amplifier (MCS, Russia). Passive Ag/Cl sensors were placed in accordance with the 10/10 international montage system. The TP10 electrode was used as the reference. The skin-electrode impedance for each of the electrodes did not exceed 20 kΩ. Bandpass filtering was carried out in the range 0.1-75 Hz using a FIR filter. An additional filtering was conducted using a 50-Hz Notch filter. Raw data collection was carried out using the NeoRec software (MCS, Russia) synced with the stimulus presentation environment. Stimulus presentation was carried out via a self-written code based on the PsychoPy python-module [44]. We also used one bipolar lead to record EMG from the *flexor digitorum superficialis* (FDS).

### Signal Preprocessing

Raw EEG recordings were re-referenced using the common average reference (CAR), which also has spatial filtering effects [45]. Next, the signals were band-pass filtered in the range 1-30 Hz using a fourth-order Butterworth filter. Noisy channels were interpolated using the spherical spline method [46]. The preprocessed signal was divided into 8-s epochs (from -1 to 7s relative to trial onset) for each trial. To assess the time-frequency dynamics of EEG oscillations amplitude, the Morlet wavelet transform with variable number of cycles was applied for all the extracted epochs. The frequencies of the wavelets ranged from 6 Hz to 30 Hz with 0.4 Hz step. The full width at half maximum (FWHM) was equal to 180 ms corresponding to a spectral FWHM of 6Hz. Then the time-frequency matrices with Morlet coefficients were converted to ERD values with Eq.1, where Morlet coefficients for each timestamp within the trial were used instead of median PSD value.

To plot the topographic distribution of ERD/S in each experimental condition, the power spectral density was calculated by the Welch method [47] for 5 sec epochs (1-6s) since the trial onset. The subject-specific peak frequency was selected semi-automatically (i.e., under visual inspection) within the range 8-15 Hz. The subject-specific frequency subrange was selected around the peak frequency ± 1.5 Hz and then was used for ERD calculation in each particular channel position:

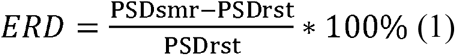

Where PSDsmr is the matrix power spectral density median value of all the TS, TI or control trials in the range 1-5 sec and subject-specific frequency subrange for each channel position. PSDrst defines the same size matrix of PSD calculated in the same manner for trials corresponding to the resting state.

### Statistical Analysis

To determine significant changes in the time-frequency dynamics of oscillatory activity and spatial distribution of the ERD/S values in all the experimental conditions (TS, TI and control), the non-parametric cluster-level paired t-test with 10.000 permutations was used [48]. This procedure is assumption free and is widely used for multichannel data and time-frequency analysis to avoid the multiple comparisons problem. The Friedman’s test and a Wilcoxon signed ranks test as the post hoc statistical test were performed to determine the statistically significant differences in total ERD/S value within the full trial in TS, TI and control condition, as well. The Bonferroni correction was applied for the number of tested hypotheses (adjusted to p<0.003 for 15 comparisons).

Statistical analysis was performed via the SciPy module v.1.4.1 [49]. For signal processing and permutation methods, the MNE-Python v.0.23 was used [50]. For data visualization, we used the Matplotlib graphics environment [51].

## RESULTS

### Time frequency EEG dynamics during tactile stimulation and imagery

We started with the analysis of the changes in EEG rhythms for the TS and TI trials. The Wavelet-Morlet transform was applied to the signal from the C3 channel (i.e. the sensorimotor area contralateral to the stimulated hand) and used this signal to analyze the event related time-frequency perturbations (TFP). With the Morlet transformation we obtained the time-frequency representation matrices for the TS, TI and control conditions. Then, we baselined them to the resting state (data from visual-attention trials), calculated the median over the group and then visualized. Figure 2 shows that the group-average temporal spectral dynamics were similar during the TS and TI trials: the *μ*-rhythm amplitude gradually decreases relative to resting state and remains at the reduced level until the end of the trial (Fig. 2B). Then, it returned to the baseline. To discover significant differences between the somatosensory conditions (TS and TI) and the control state, we performed intra-subject subtraction of the time-frequency matrix corresponding to the control condition from the time-frequency matrices corresponding to TS and TI conditions. The non-parametric cluster-level paired t-test was used to compare the obtained difference matrices with zero. We also used a similar procedure to perform comparison between the TS and TI conditions. Both the tactile stimulation and tactile imagery were characterized by a decrease across the α and *β*-band power compared to the control condition. However, only the the decrease in the *μ*-rhythm power was significant for the TI-condition when it was compared to the control state (e.g., the *μ*-ERD clusters that persists for the entire trial duration in Fig. 2A), whereas a significant *β*-ERD occurred during the TS trials. No differences in TFP between the TS and TI were observed. Thus, the frequency range of the *μ*-rhythm response appearing was similar for TS and TI (∼8-14 Hz as can be defined on Fig. 2).

**Fig 2.**
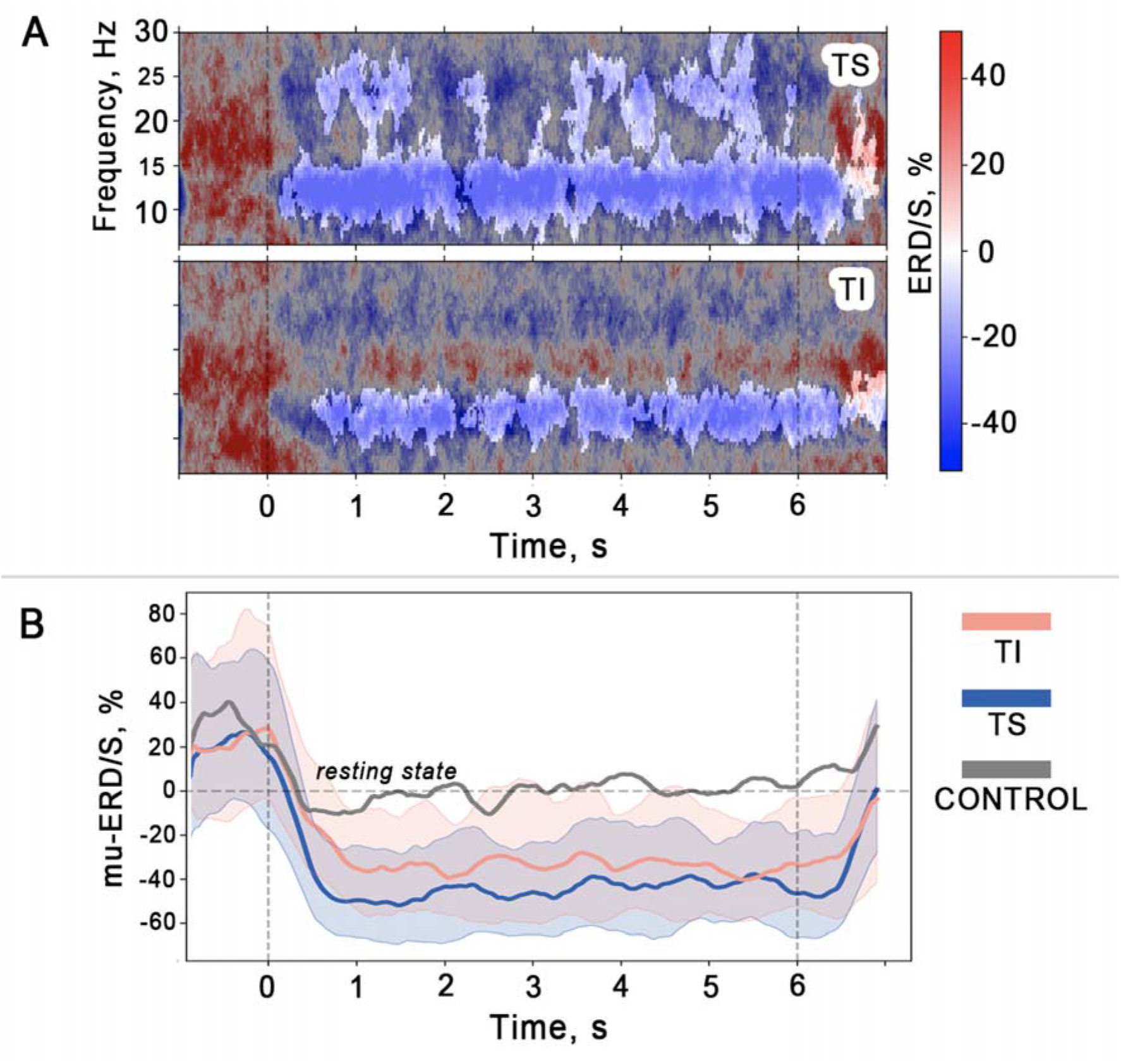
**A** - Grand median (N=17) time–frequency ERD/S distribution in the C3 channel during the tactile stimulation (TS), tactile imagery (TI) and difference (diff., TS-TI). Blue shapes correspond to the level of desynchronization (ERD), red – synchronization (ERS). The gray mask indicates insignificant differences. *(p >* .*003, non-parametric cluster-level paired t-test, p-level adjusted by Bonferroni correction)*. **B** - Grand median of the ERD/S time courses during tactile stimulation (TS), tactile imagery (TI) and control state. Color lines correspond to median values (median values were counted within each subject over all trials and individual frequency ranges where ERD was detected, and then a grand median value was found). Color shapes show the corresponding 25 and 75 percentiles. Vertical dashed lines limit 6s trial, horizontal line defines resting state.

### ERD/S topographical patterns

To determine the topographical localization of the sensorimotor response, we compared the *μ-*and *β-*ERD values for each electrode position across all experimental conditions using the nonparametric permutation tests. Median (N=17) topographic distribution patterns of *μ-*ERD/S for all experimental conditions are shown in Fig 3A. The *μ-*ERD induced by the tactile stimulation of the right hand wrist was more prominent in the central EEG sites over the sensorimotor cortical areas, with some contralateral dominance. The TI condition had a similar *μ*-ERD distribution pattern. A different pattern was found for the control state.

**Fig 3.**
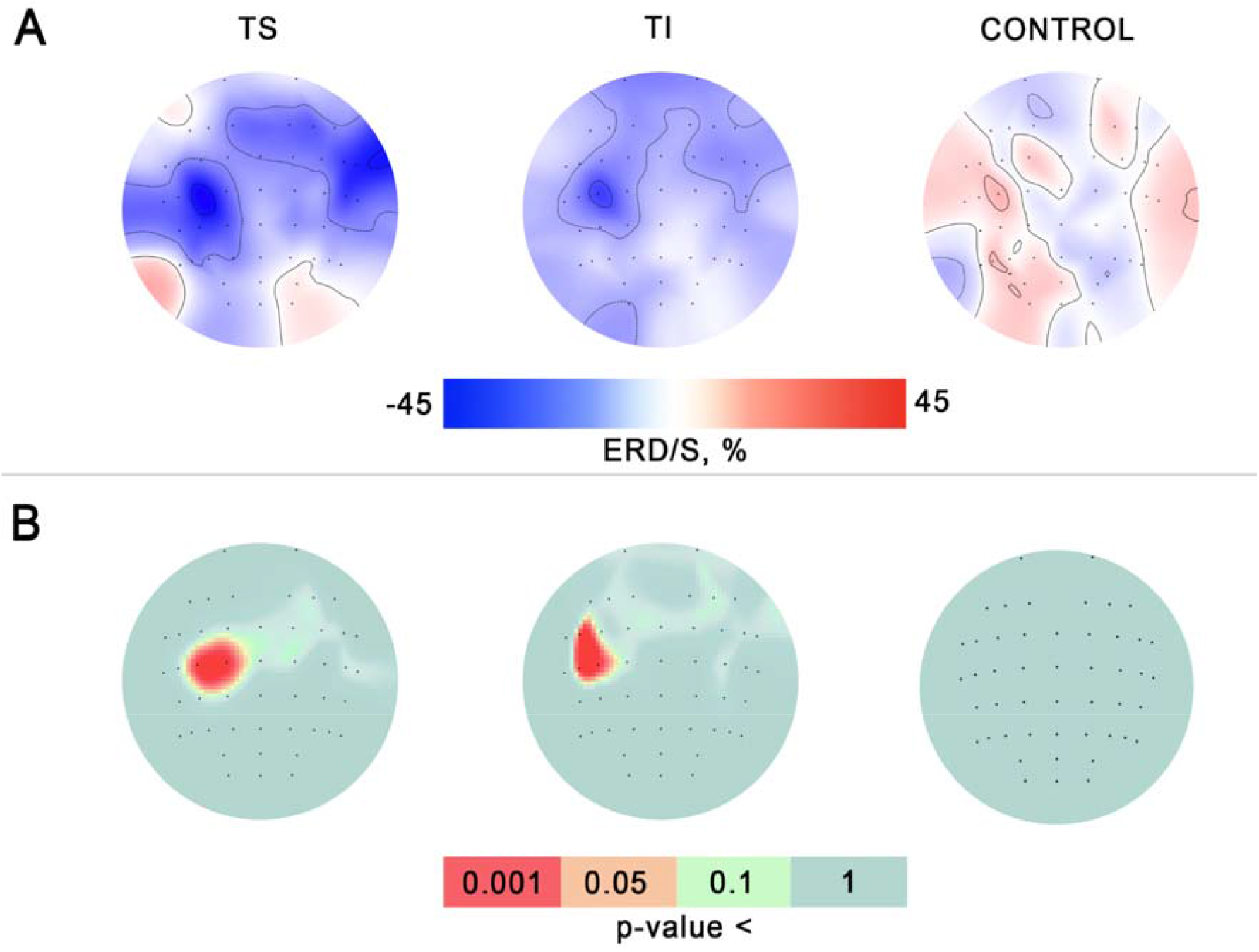
**A** - Group median ERD/S maps of sensorimotor EEG **μ-**rhythmic activity in two active conditions and the control for all subjects (N=17). Blue corresponds to the level of desynchronization (ERD), red – synchronization (ERS). **B** - The across-subjects significance maps visualized p-values derived from non-parametric permutation tests for all experimental conditions. Color defines the significance level. Red color marks electrodes in which there was across-subjects significance.

Using a permutation t-test relative to zero for each experimental condition, we determined the channels where significant *μ*-ERD occurred. The permutation t-test revealed a significant decrease in the μ-rhythm amplitude over C3 and C1 channels for the TS condition (*tC3 = - 5*.*9; p = 0*.*001; tC1 = -6*.*5; p = 0*.*0007*) and over FC5, C3 and C5 for the TI condition (*tFC5= -6*.*2; p = 0*.*0009; tC3 = -5*.*9; p = 0*.*0015; tC5 = -7*.*3; p = 0*.*0002*). No significant *μ-*ERD topographical origins were found in the control state, that is when the tactile stimulation was not applied and participants did not perform imagery of tactile sensations. The obtained matrices of p-values were superimposed over the scalp EEG channels positions for visualization (see Fig. 3B). Since the TS-trials were characterized by a significant event-related desynchronization in the ∼20-26 Hz frequency range, we performed spatial permutation t-test for the *β-*ERD values.

Figure 4 shows that a significant decrease in *β-*amplitude induced by the tactile stimulation of the right-hand wrist occurred contralaterally, predominantly over the fronto-central EEG channels. Significant spatial clusters where *β-*ERD emerged were discovered neither in the TI condition nor in the control state.

**Fig 4.**
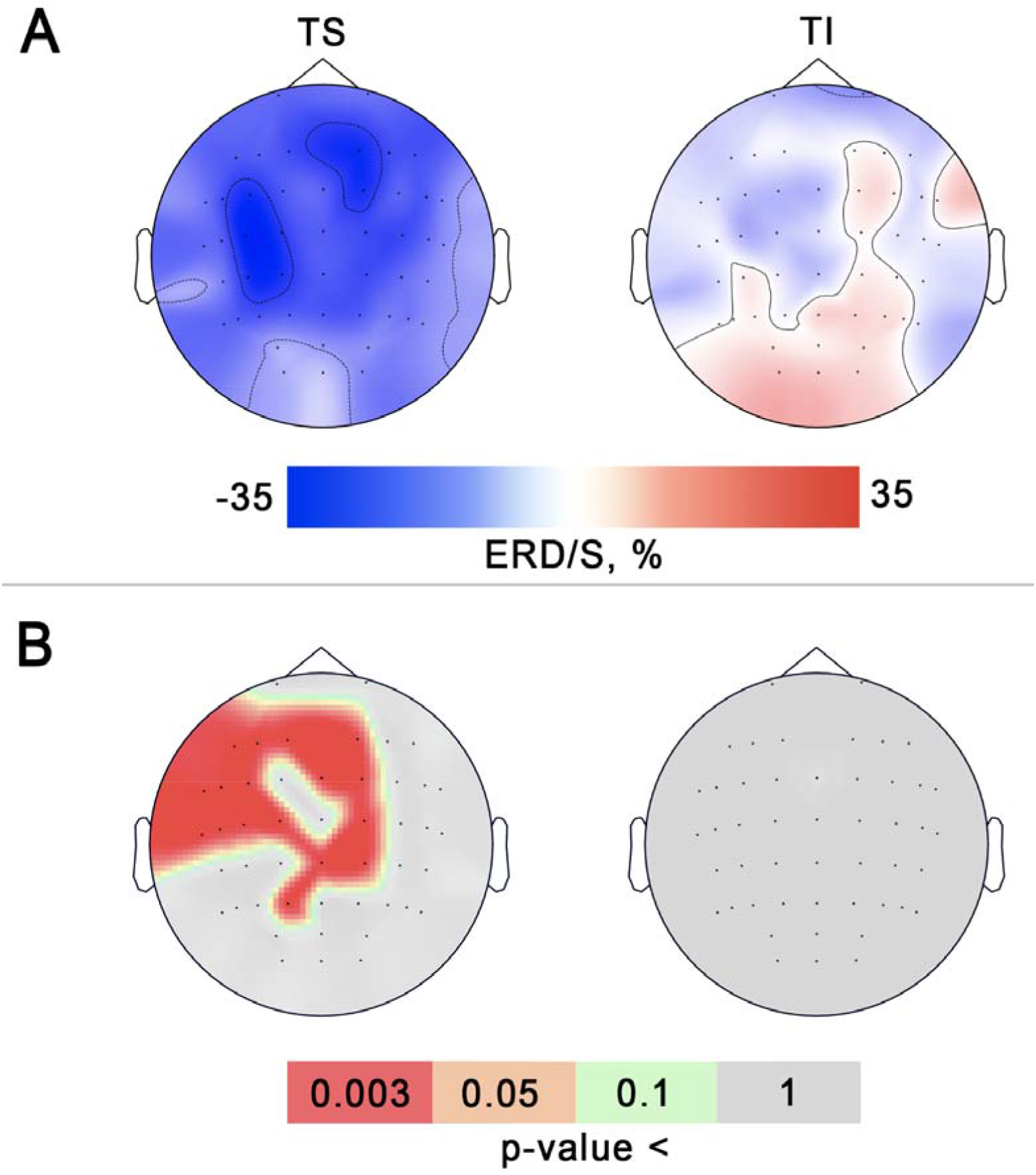
**A** - Median ERD/S maps of sensorimotor EEG β-rhythmic activity in two active conditions for all subjects (N=17). Blue corresponds to the level of desynchronization (ERD), red – synchronization (ERS). **B** - The across-subjects significance maps visualized p-values derived from non-parametric permutation tests for all experimental conditions. Color defines the significance level. Red color marks electrodes in which there was across-subjects significance.

To analyze the global *μ-* and *β-*amplitude decrease within the epoch duration, we calculated the median ERD value over all trials for the interval starting with the 1st second till the end of the trial, and over the individual frequency ranges where ERD was detected across all the participants. We rejected the 1st second of each experimental trial because of the TI onset jitter, which contributed to a gradually developing ERD (see Fig. 4). The 1st second of the visual-attention trials was rejected to avoid the residual effects of the tactile stimulation and imagery. There were significant differences between the experimental conditions according to Friedman’s test in *μ-*ERD value in the C3 channel (*χ2=25*.*7; p = 0*.*000002*). The paired comparisons revealed no significant difference between the ERD values in TS and TI conditions (*W=56; p=0*.*33*; see Fig.5). On the other hand, as shown in Fig. 5, the *μ-*desynchronization in TS and TI conditions was significantly stronger compared to the control state (*W=0; p < 0*.*0001*).

**Fig 5.**
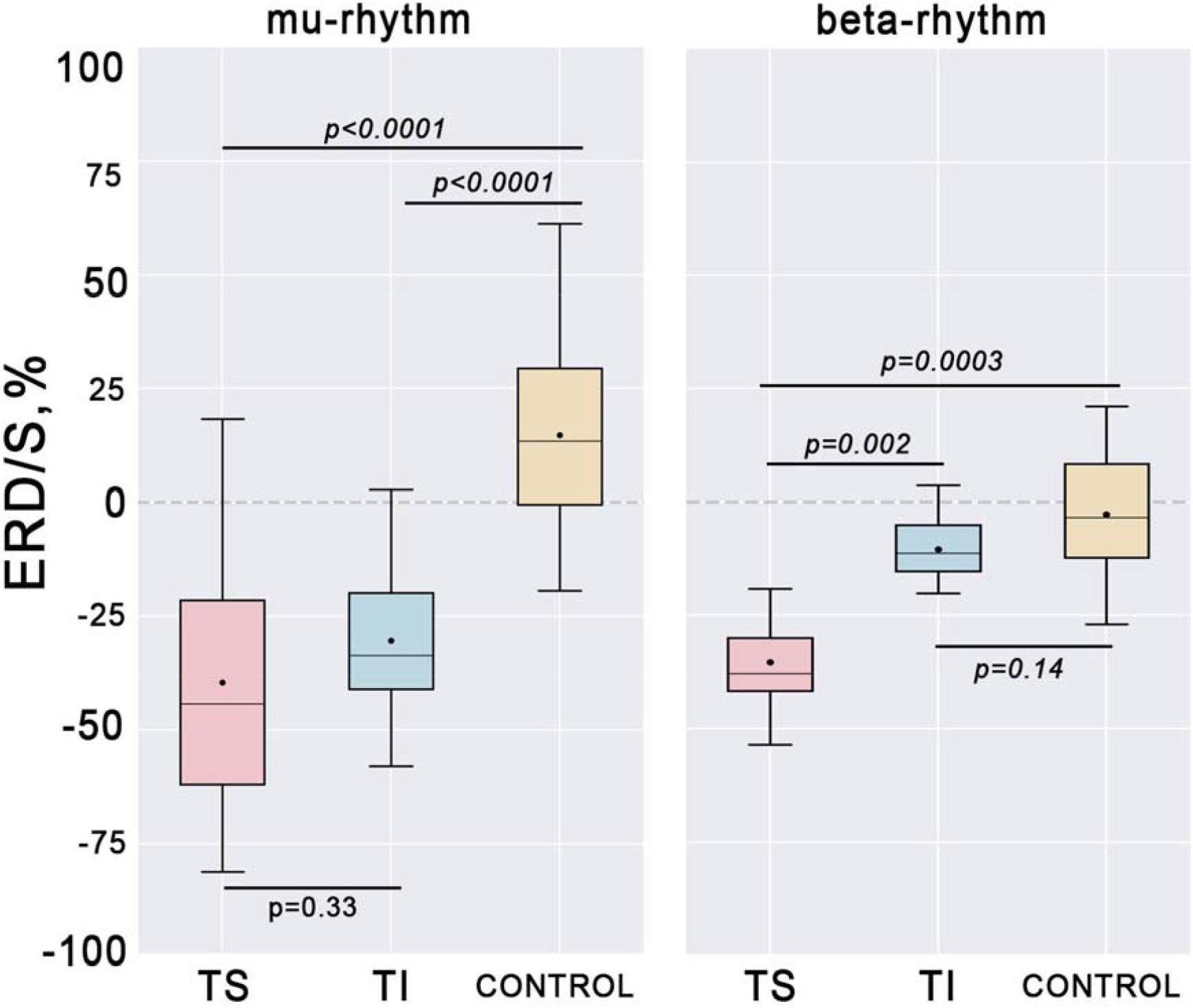
Group level (N=16) ERD-values in the C3 channel for explored somatosensory conditions (tactile stimulation, tactile imagery, and control condition). Horizontal lines within the boxes - medians, boxes – interquartile range and [Q1-1.5*IQR; Q3+1.5*IQR] range is shown by whiskers. Circles represent mean values for the group. P values are shown for the Wilcoxon signed-rank test.

A statistical analysis of the β-ERD values revealed significant differences between the three experimental conditions (*χ2=15*.*6; p = 0*.*0004*). Further analysis (Wilcoxon tests) indicated that *β-*ERD was significantly stronger in the TS-condition as compared to the and Control conditions (*W=13; p = 0*.*002; W=1; p=0*.*0003*, respectively).

## DISCUSSION

In this study, we examined the changes in sensorimotor rhythms during real and imagined vibrotactile stimulation. We found that vibrotactile stimulation of the wrist as well as imagery of vibration-induced sensations caused a prolonged decrease in the EEG power in the *μ-*range. Time-frequency analysis showed a significant *μ-*ERD in the 9-14 Hz range throughout the 6-s TS or TI trials. Additionally, TS (but not TI) caused significant β-ERD in the frequency range of 20-26 Hz. The frequency range where sensorimotor ERD appeared during TS and TI, as well as its spatial distribution, were similar to the well-known ERD patterns associated with real and imaginary motor activities.

### Sensorimotor ERD during tactile stimulation and tactile imagery

Electrical stimulation of the median nerve is a widely used method for exploring the cortical effects of afferent stimulation. Many studies showed that median nerve stimulation leads to prominent contralateral ERD of the sensorimotor EEG rhythms [52, 53]. Similar results were observed in the studies with cutaneous vibro-tactile stimulation [24, 54]. Here we used vibrotactile stimulation of the skin area on the right wrist. We expected to obtain results consonant with the results of aforementioned studies. Such effects of afferent stimulation suggest a general principle for the relationship between α-like rhythms and sensory events: prior to the presentation of somatosensory stimuli, synchronized α activity is observed, and the stimuli result in α-desynchronization, possibly because of disinhibition of thalamocortical circuits [55].

In addition to *μ-*ERD, tactile stimulation led to the desynchronization in the *β*-frequency range. Yet, this effect was not observed for TI (i.e. was statistically insignificant). *β*-oscillations have been linked to the activity of the primary motor cortex activity as it is well known that somatosensory inputs modulate *β* oscillations because of motor-cortical mechanisms [56-58]. These effects are driven by the ascending projections to M1 [59, 60] and by corticocortical connections with the somatosensory cortex [61]. We suggest several reasons of absence of the β-ERD during TI. The first possibility is that TI is maintained by sustained activity of the primary and secondary somatosensory areas without the engagement of the motor cortex [28, 62]. Indeed, participants did not imagine any movements in our experiments, so the activation of motor circuits could be weak. The second possibility is that the motor cortex responded during TS but not TI is that application of vibration to FDS generated a strong input to the motor cortex because of the activation of muscle spindle afferents [63]. Stimulation of muscle spindles could evoke kinesthetic illusions, but this was not the case in our experiments particularly because the participants were instructed not to imagine limb movements. The third possibility is a relatively low level of alertness during TI compared to TS. Thus, Ilman et al., 2021 suggested that alertness level has an effect on the sensorimotor *β*-desynchronization during tactile stimulation. Yet, the effects they observed were not statistically significant at the group level [57].

Notably, despite the absence of statistically significant β ERD during TI, we observed a slight decrease in β power in this condition. This matches the general mental imagery model, which considers mental imagery as a reverse form of perception process with weaker cortical manifestations [64-66]. According to this model, mental imagery is based on the processing of information retrieved from memory by cortical areas. Following this idea, we propose that tactile stimulation can lead to memory formation, and this memory could be brought back to somatosensory areas when the previously experienced sensation is reproduced by imagining. Our results and the proposed interpretation agree with the simulation theory of motor imagery formulated by Jeannerod [67].

### Tactile imagery perspectives

The tactile sense is one component of the complex motor image, in fact, the tactile imagery is a part of the motor imagery, which includes kinesthetic sensations and visual image as well. Motor imagery is a widely used mental technique in sport to improve performance and motor imagery based BCIs are promising tools for the post-stroke motor recovery. However post-stroke patients often have difficulties in motor imagery learning. Tactile imagery similarly to motor imagery can be proposed as a BCI control paradigm based on *μ*-ERD detection as well [68]. We suppose that somatosensory mental images are easier to perform and the learning of tactile imagery is more regulated and understandable compared to motor imagery, thus TI-based BCIs could be more useful and lower barriers to entry compared to MI-BCIs. There are a lot of studies showing the usefulness of tactile feedback and tactile stimulation during motor imagery training and MI-BCI control [69-71]. We believe that the positive effect of TS on the learning on kinesthetic imagination of movements may be associated with the use of mental tactile sensations (i.e. participants can actually use the tactile imagery to increase the vividness of motor image and induсe the sensorimotor rhythms depression more effectively).

### Significance of the obtained results

While TI was the subject of several previous studies, here we studied its manifestation in EEG for the first time. We found μ-rhythm desynchronization in the contralateral hemisphere during both TS and TI and β-desynchronization during TS only, which we interpreted as activation of both somatosensory and motor cortices during TS and TI and activation of somatosensory cortex only during TI. Our results generally match the plentiful previous results on motor imagery assessed with EEG technique [72, 73]. The results we obtained also match the fMRI studies where tactile imagery was found to increase somatosensory cortical activity and enhance functional connectivity between the prefrontal and somatosensory cortical areas [29, 31]. The other study that our results agree with is the work [32] where microelectrode techniques were used to localize cortical regions involved in the tactile image formation. Our work is also relevant to the research of spinal circuits where tactile and kinesthetic types of processing lead to increased spinal excitability [74]. Overall, our work adds to the existing literature on TI and is also relevant to the field of brain-computer interfaces (BCIs), where TI could be used to obtain EEG responses desired for communication, control of external devices and rehabilitation purposes.

### Limitations

The results of the current study provide convincing evidence that TI leads to contralateral _*μ*_-rhythm desynchronization, and this effect is similar to the one evoked by the real vibrotactile stimulation applied to the wrist. However it is important to mention several limitations to our research. First, EEG recordings allowed us to r**eated** ERD/S to sensorimotor responses, but we could not localize the source of ERD precisely with this method. High-density EEG recordings combined with inverse solution computing (i.g. MNE, sLORETA) could provide more detailed information about the sources of TI-related activity and could help to better compare the effects of TS and TI. Other neurophysiological techniques besides EEG, like TMS, MEG, fNIRS, could be useful for this purpose, as well.

We used a visual scene as a reference state and also used it in the control condition to exclude the conditioning effect of the visual pictogram used for the TS/TI tasks. The obtained results suggest that sensorimotor rhythm modulation was the specific effect of TI. Yet it could be useful to have an additional control for tactile attention in future studies. There could be a somatosensory attention condition, where the subject’s attention is drawn to the stimulation site, but without performing imagery.

Another limitation in this study is participants’ age range since most of the subjects were relatively young (from 19 to 30 years old). It is important to complement our findings with a broader-age group in the future. Since a decline in sensorimotor functions is typically observed with aging, it would be of interest to study the effect of age on the EEG patterns.

## CONCLUSION

Based on EEG recordings conducted during tactile imagery, we conclude that the effects of TI (namely, *μ*-rhythm desynchronization) are similar to the effects of MI. Such mutuality of motor and somatosensory cortical processing, at least at the level of EEG, could indicate that the “imagery circuit” of the brain is not strictly localized. Yet, by changing imagery requirements it is possible to change activity distribution in somatosensory versus motor areas. Considering the sensory origin of the α rhythms in general, α-desynchronization during TI is yet another phenomenon where somatosensory processing is affected by thought. As such, it could be used in BCI design where thought needs to be communicated from the brain to the external devices.

## Author Contributions

LY proposed the general concept of the study. Experimental design was developed by LY, NS and AK. The experimental sessions were performed by LY, NS and AM. Data analysis and visualization was performed by LY and NS. LY wrote the first draft of the manuscript. LY, NS, AK and ML revised the manuscript. All authors contributed to the final revision, read, and approved the submitted version.

## Acknowledgments

The work of L. Yakovlev, N. Syrov and A. Kaplan was supported by the Russian Science Foundation under grant #21-75-30024. The work of A. Miroshnikov was performed within the scope of the State Assignment of the Ministry of Education and Science of the Russian Federation #FZWM-2020-0013 and the Priority 2030 Program.

